# Iterative delivery of mRNA by MEFI

**DOI:** 10.1101/2025.07.31.667827

**Authors:** Yun Gao, Huan Li, Yue Xun, Yu Liu, Yaoyang Zhang, Shuai Liu, Xiaobin Fang, Xiaojiao Shan, Junhua Liu, Qinyao Zhu, Xiushan Yin, Luo Zhang

**Author notes:** Correspondence (X.Y.), (L.Z.).

## Abstract

Conventional therapeutic mRNA delivery systems are inherently single-use, limiting expression potency, duration, and penetration across biological barriers. To address this, we engineered the mRNA Exporting and Ferrying Implement (MEFI), a platform that enables iterative cell-to-cell transfer of cargo mRNA, allowing a single mRNA molecule to be utilized multiple times across various cells. MEFI encodes a fusion protein that assembles virus-like particles (VLPs) and encapsulates both itself and cargo mRNA into these VLPs, orchestrating reproduction of VLPs from targeted cells and continuous intercellular spreading. MEFI significantly amplified cargo protein expression in the transfected cell populations (∼20-fold) and in organs (∼500-fold in lung) from mice administered systemically. Notably, through intramuscular injection of naked plasmids, MEFI promoted robust (∼20-fold) and durable (∼2-month) cargo expression, interorgan mRNA transfer, and successful crossing of the blood-brain barrier. Moreover, modified MEFI demonstrated cell-type-specific targeting, mediating the elimination of antigen-expressing cells. Collectively, MEFI establishes a modular platform for enhancing mRNA delivery, leveraging the concept of iterative delivery of mRNA.

**Graphical Abstract:** 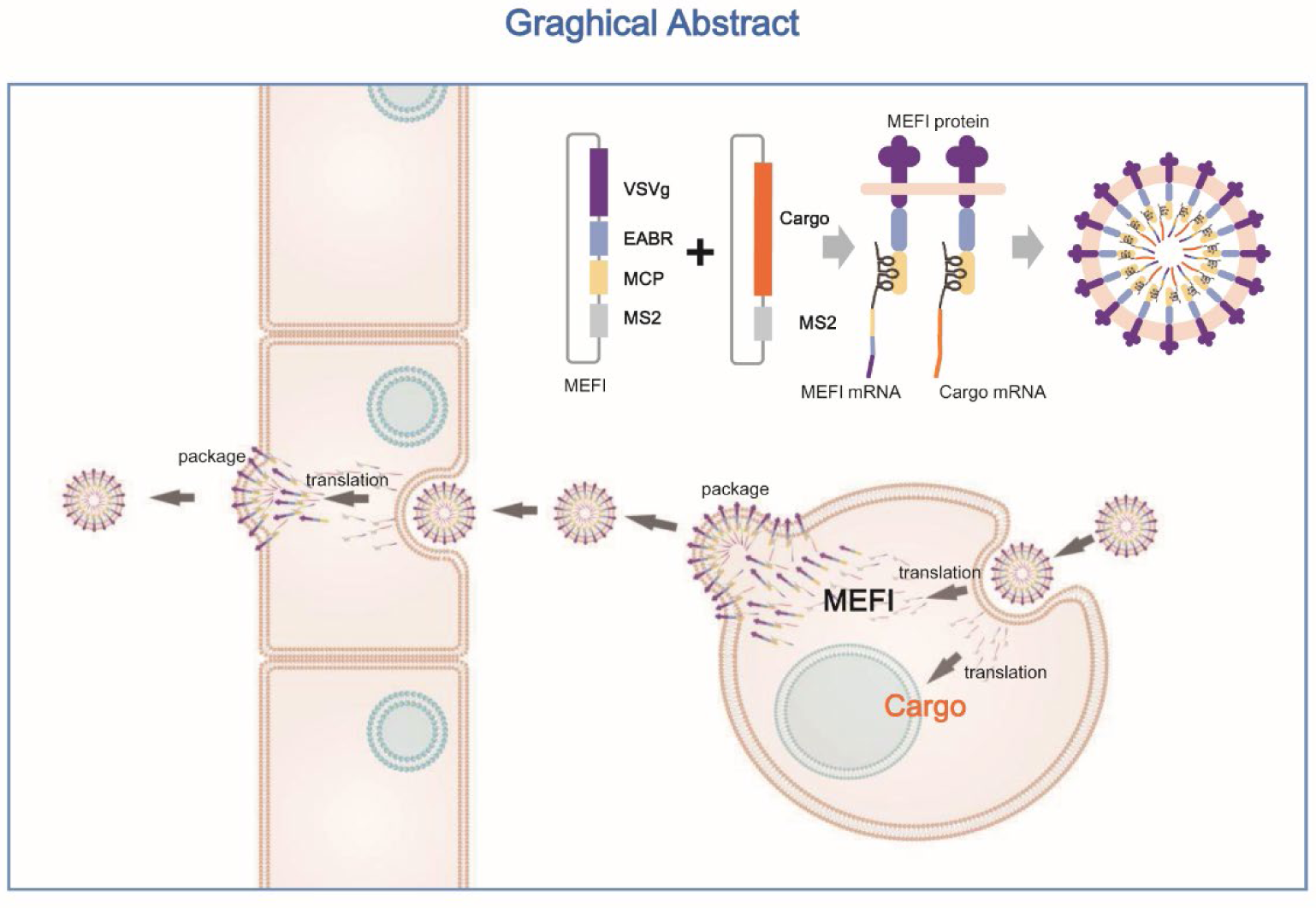

## Introduction

Messenger RNA (mRNA)-based therapeutics have emerged as a transformative platform for treating diverse diseases, including infectious diseases, cancers, and genetic disorders^1–4^. Innovations in mRNA engineering and lipid nanoparticle (LNP) delivery have significantly enhanced stability and translational efficiency, exemplified by the rapid development of COVID-19 vaccines^5,6^. However, a fundamental limitation persists: current mRNA therapies operate through a "single-use" paradigm - delivered mRNA functions transiently within individual cells before degradation. This restricts protein production, hampers delivery across biological barriers (e.g., blood-brain barrier), and limits efficacy in poorly accessible tissues. To overcome these constraints, strategies enabling intercellular mRNA transfer are highly desirable.

Virus-like particles (VLPs) present a promising approach for the transfer of intercellular mRNA by encapsulating cargo during the budding process from producer cells^7–14^. Yet, conventional VLP systems require multiple viral components (e.g., gag, pol) for assembly^15–20^, increasing complexity and immunogenicity^21^. While the overexpression of the vesicular stomatitis virus glycoprotein (VSVg) can lead to the release of simplified VLPs, their packaging efficiency for cargo mRNA remains low in the absence of capsid proteins^22^. Critically, no existing system achieves iterative transfer, where cargo mRNA delivered to primary cells initiates secondary production of VLPs, amplifying cargo expression across cell populations.

Here, we propose a minimalist design to enable self-propagating mRNA delivery. Many enveloped viruses recruit endosomal sorting complex required for transport (ESCRT)-associated proteins such as TSG101 and/or ALIX through capsid to conduct the budding and releasing process^23–29^. TSG101 and ALIX are also recruited by the protein CEP55 through a short amino acid sequence termed ESCRT- and ALIX-binding region (EABR) in the process of cytokinesis^30–32^. Thus, fusing the EABR to the cytoplasmic tail of a viral glycoprotein or other membrane protein directly recruits TSG101 and ALIX, bypassing the need for co-expression of other viral proteins for VLP self-assembly^33^. The aptamer MS2 and its interacting aptamer-binding protein, MS2 coat protein (MCP), have been widely used^34–36^. VSVg mediates broad tropism and fusion via LDL receptor interaction^37,38^. By integrating these elements, we engineered a single chimeric protein, VSVg-EABR-MCP (VEM), that autonomously assembles mRNA-packaging VLPs. Furthermore, we inserted MS2 aptamers into the 3’UTR of VEM mRNA, creating a self-packaging entity termed mRNA Exporting and Ferrying Implement (MEFI). This design enables MEFI to: export both cargo mRNA and itself via VLPs; initiate VLP re-production in target cells, establishing iterative transfer cycles; serve as a modular platform capable of cell-specific delivery. Collectively, MEFI establishes a paradigm for self-propagating mRNA delivery via recursive VLP assembly and intercellular transfer.

## Results

### VSVg-EABR-MCP fusion protein assembles VLP that enabling MS2-tagged mRNA delivery

Traditional VLPs typically rely on capsid proteins for self-assembly and fusogens for target-cell binding and fusion. To engineer a single protein fulfilling both roles, we fused the human CEP55-derived EABR domain to the C terminus of VSVg’s cytoplasmic tail via a flexible Gly-Gly-Gly-Ser linker, appending green fluorescent protein (GFP) for visualization (Figure 1A). The transient transfection of the VSVg-EABR-GFP plasmid led to the localization of GFP around the membrane in human HEK293T cells (Figure S1A). Electron microscopy images indicated that spherical, ∼160-nm-diameter particles were contained in the supernatant (Figure. 1B). Subsequently, the supernatant was collected and concentrated to infect murine RAW264.7 cells. Fluorescence microscopy and flow cytometry assays indicated that the majority of the cells were GFP-positive, suggesting that the VSVg-EABR-GFP VLPs mediated effective cross-species transduction (Figure 1C and Figure S1B).

**Figure 1.**
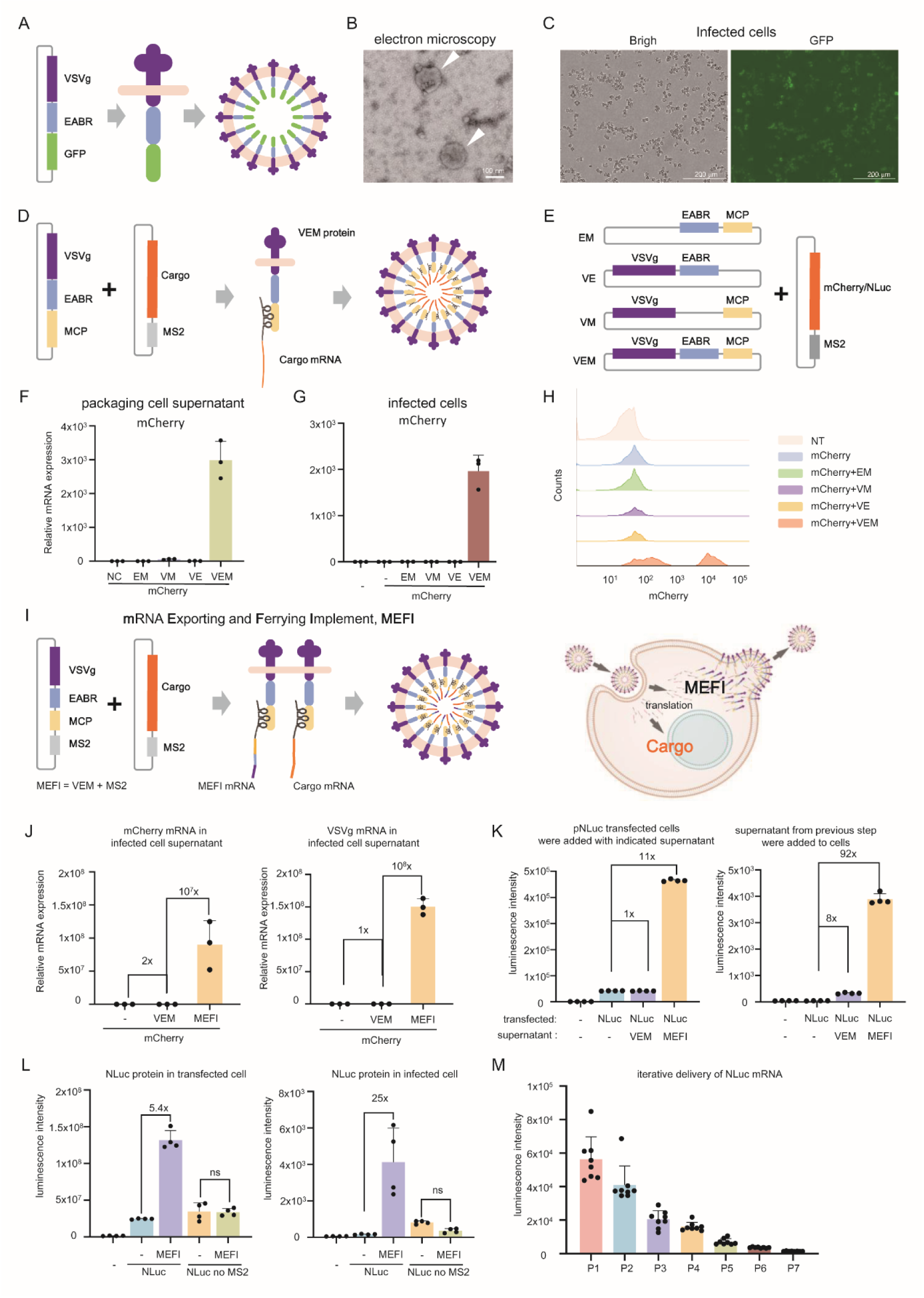
MEFI encodes fusion protein packages mRNA into self-propagating VLPs, enabling continuous cell-to-cell transfer. (A) VSVg-EABR-GFP fusion protein facilitates the assembly of VLP (schematic). (B) VSVg-EABR-GFP VLP purified from transfected 293T cell culture supernatants by ultracentrifugation on a 20% sucrose cushion. Particles were visualized using uranyl acetate negative staining and imaged using a transmission electron microscope. Scale bars, 100 nm. Magnification, ×180,000. (C) RAW264.7 cells were infected with VLPs from B for 24 h before fluorescence images were taken. (D) The schematic diagram shows VSVg-EABR-MCP fusion protein facilitates the assembly of VLP, which packages cargo mRNA. (E) The schematic diagram illustrates the composition of VSVg-EABR-MCP and its truncations. (F) The relative quantity of mCherry-MS2 mRNA in the supernatants of 293T cells transfected for 36 h with the indicated plasmids by qPCR assay. Each dot represents one technical replicate. (G) The relative quantity of mCherry-MS2 mRNA in RAW264.7 cells infected with the supernatants indicated in (F) for 24 hours by qPCR assay. Each dot represents one technical replicate. (H) Flow cytometry conducted on cells from (G). (I) The schematic diagram shows MEFI facilitates the assembly of VLP, which packages both cargo and itself mRNA, enabling reproduction in infected cell. (J) The indicated plasmids were transfected into 293T cell. After 36 hours, the supernatants were filtered and added to RAW264.7 cells. Then, the mixed culture media was replaced with fresh media 6 hour later to remove residue VLPs, and 24 hr later, the supernatants were collected for qPCR assay. (K) RAW264.7 cells were transfected with pNLuc plasmids, and 293T cells were transfected with either VEM or MEFI plasmids. After 24 hours, the supernatants from the transfected 293T cells were added to the RAW264.7 cells. The expression of NLuc in these cells was then detected using chemiluminescence after an additional 24 hours (left). Concurrently, the supernatants from the RAW264.7 cells were transferred to a fresh batch of RAW264.7 cells. 24 hours later, the expression of NLuc in these cells was also measured using chemiluminescence (right). (L) The indicated plasmids were transfected into 293T cells. After 36 hours, the NLuc expression was measured using chemiluminescence (left). Concurrently, the supernatants were transferred to RAW264.7 cells. Another 24 hours later, the expression of NLuc in these cells was measured using chemiluminescence (right). (M) 293T cells were co-transfected with NLuc-MS2 and MEFI plasmids, 36 hours later, the supernatant were filtered and added to MEFs. After 6 hours, the media was replaced with fresh one. Another 18 hours later, the expression of NLuc was measured, and the supernatants were transferred to a fresh batch of MEFs. This round was conducted 7 times as described in Fig S1D.

For mRNA delivery, GFP was replaced with the MS2 coat protein (MCP) to recruit MS2-tagged cargo mRNAs (Figure 1D). To define the roles of VSVg, EABR, and MCP in mRNA packaging and delivery, we generated element-deletion variants (Figure 1E). Co-transfection of mCherry-MS2 with the VSVg-EABR-MCP chimera (VEM) resulted in >2,000-fold enrichment of packaged mCherry-MS2 mRNA in supernatants of HEK293T cells (RT-qPCR), relative to controls lacking VEM. In parallel, we observed that VSVg, EABR, and MCP were all essential for mRNA packaging (Figure 1F). Subsequently, the filtered supernatant was used to infect RAW264.7 cells, and the cargo mRNA in these cells was assayed. As expected, VEM efficiently delivered mCherry mRNA into target cells. In contrast, negligible cargo mRNA was detected when either VSVg, EABR, or MCP was omitted (Figure 1G). Similar results were obtained when NLuc was used as the cargo (Figure S1C). In agreement, the expression of mCherry protein in infected cells by flow cytometry assay confirmed that all three elements are necessary for functional mRNA packaging and delivery (Figure 1H). These results suggest that VSVg-EABR-MCP fusion protein assembles VLP that enabling MS2-tagged mRNA package and delivery.

### MEFI orchestrates cargo mRNA transfer from cell to cell iteratively in vitro

Current mRNA delivery systems confine translation to a single cell before degradation. We posited that the iterative production of VLPs packaging cargo and VEM mRNA from targeted cells could transfer these mRNA to other cells and amplify the protein output of the cell population. To enable this, we engineered MEFI by inserting MS2 aptamers into the 3′UTR of VEM mRNA, enabling self-packaging into VLPs upon translation (Figure 1I).

To determine the efficiency of MEFI to re-package and export cargo mRNA, mCherry alone, mCherry plus VEM, and mCherry plus MEFI plasmids were transfected into 293T cell, respectively. After 36 hours, the supernatants were filtered and added to the RAW264.7 cells. Then, the mixed culture media was replaced with fresh media 6 hours later to remove residue VLPs, and 24 hours later, the supernatants were collected for qPCR assay. The results indicated that MEFI enriched almost 10^8^-fold of mCherry mRNA compared to mCherry alone. In contrast, VEM enriched mCherry mRNA by ∼2-fold compared to mCherry alone. In parallel, a similar result was obteined when assessing the VSVg mRNA, suggesting MEFI protein can efficiently export itself mRNA (Figure 1J).

To determine whether external MEFI can export endogenous MS2-tagged mRNA from targeted cells, we transfected the NLuc-MS2 plasmid alone, which is convenient for quantitative detection, into RAW264.7 cells. After 24 hours, the indicated supernatants were added, and NLuc in the cells was detected by chemiluminescence following an additional 24-hour infection period. We observed that the addition of VEM supernatant did not increase the expression of NLuc, while MEFI increased the expression of NLuc by approximately 10-fold. Subsequently, the cell supernatants from the previous step were added to fresh RAW264.7 cells. The luminescence assay revealed that the supernatant produced by VEM exhibited a slight increase, whereas MEFI achieved an approximate 100-fold enhancement in NLuc expression, thereby demonstrating intercelluar ferrying of cargo mRNA (Figure 1K). We then showed that MS2 is indispensable for cargo mRNA intercellular transfer, as MEFI does not enhance the expression of NLuc mRNA without MS2 in cells by transient plasimd transfection and supernatant infection (Figure 1L). Next, in order to assess the efficiency of multiple rounds, or iterative, delivery of mRNA by MEFI, mouse embryo fibroblasts (MEFs) were used to infect with the supernatant of previous round (Figure S1D). Although each round of Nluc expression is attenuated over the previous one, we could still detect the expression of Nluc after seven rounds of MEFI propagation (Figure 1M). Collectively, these results provide compelling evidence that MEFI can facilitate the iterative delivery of cargo mRNA through recursive VLP assembly and intercellular transfer.

### The optimized MEFI system enhances cargo expression and crosses biological barriers

Now that MEFI is capable of conducting cargo mRNA transfer from cell to cell, we next determine whether MEFI can enhance the production of total cargo protein. First, GFP-MS2 alone, GFP-MS2 plus VEM, and GFP-MS2 plus MEFI plasmids were transfected into 293T cell, respectively. The flow cytometry assay indicated that co-transfection with VEM increased the rate of GFP^+^ cells and the mean fluorescence intensity (MFI) of the total cell population compared to that of GFP alone. Furthermore, MEFI significantly increased the rate of GFP^+^ cells and the MFI of the total cell population. Additionally, we found that neither VEM nor MEFI decreased the mean fluorescence intensity (MFI) of the GFP^+^ cells, suggesting that the exporting of GFP mRNA did not reduce GFP protein production in individual cells (Figure S2 A). Moreover, the single MEFI-cargo fused mRNA did not transfer as effectively as the pattern of separated MEFI and cargo mRNA (Figure S2 B-C). Then, we performed codon and secondary structure optimization of MEFI mRNA using the LinearDesign tool^39,40^: Codon Adaption Index of MEFI increased from (CAI) 0.75 to 0.93, minimal free energy (MFE) decreased from -856.30 to -1478.40 kcal/mol (Figure S2 D). We found that the optimized MEFI had an ∼2 folds improved capacity to enhance Nluc expression (Figure S2 E). Thereafter, MEFI indicates the optimized version.

Next, we optimized the mass ratio of cargo to MEFI by transfecting a fixed quantity of the NLuc plasmid (pNLuc) with varying quantities of the MEFI plasmid (pMEFI).We observed a 5-fold increase in NLuc expression at a initial ratio of 1:0.5 compared to that at 1:0 (NLuc alone). MEFI exhibited maximal potency at a 1:2 ratio, with a 25-fold enhancement in expression compared to NLuc alone (Figure 2A). When the total plasmid quantity was fixed, the highest NLuc expression occured at a ratio of 1:2 again, with a 8-fold increase in expression compared to NLuc alone (Figure 2B). We also observed that while achieving the same level of NLuc expression, MEFI reduced total plasmid amount to at least one-tenth of that required for pNLuc alone (Figure 2C). Thereafter, unless otherwise specified, we adopt a MEFI: cargo ratio of 2:1.

**Figure 2.**
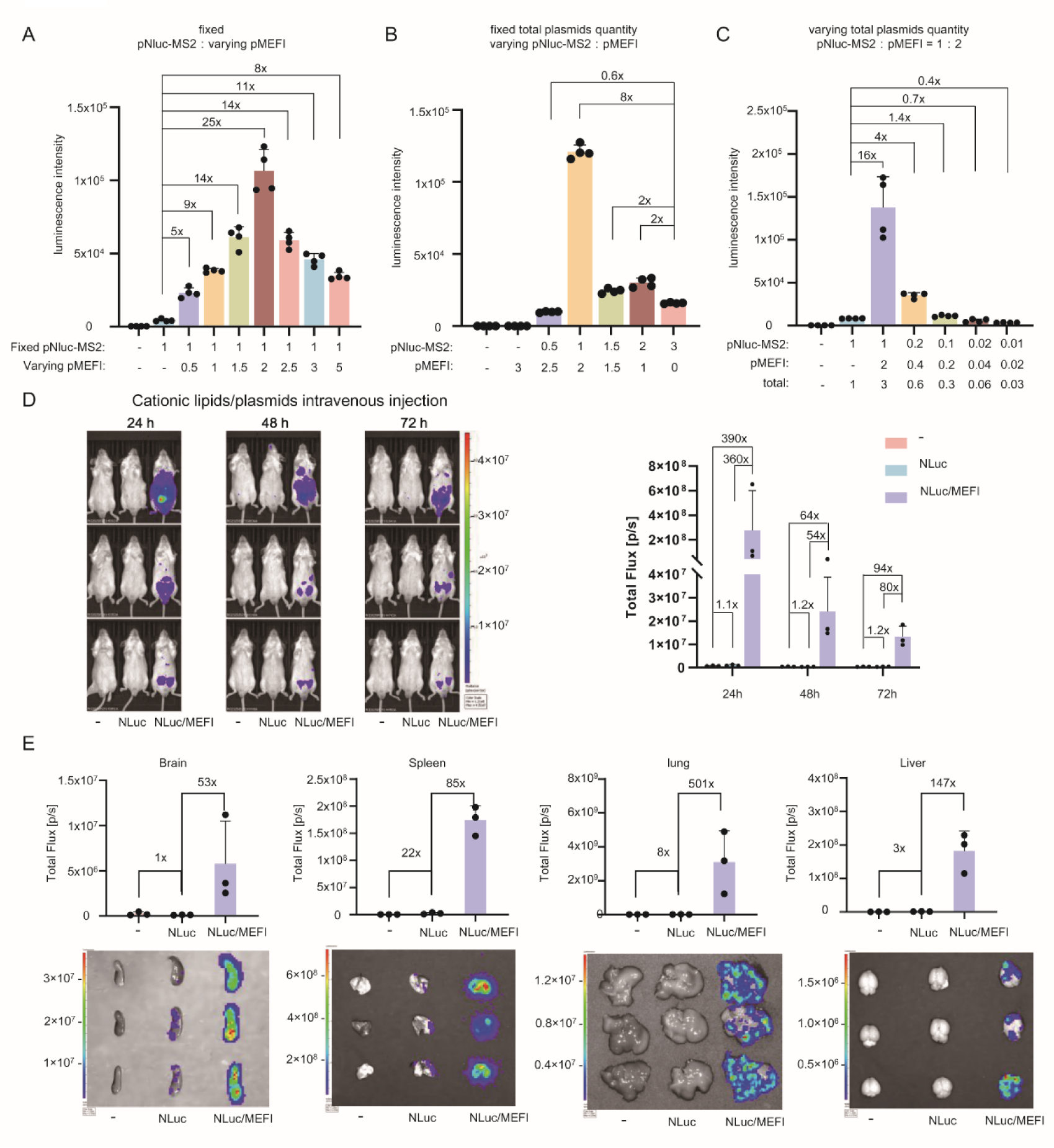
The optimized MEFI system enhances cargo expression *in vitro and in vivo*. (A-C) The indicated plasmids were transfected into RAW264.7 cells. After 36 hours, the NLuc expression was measured using chemiluminescence. (D) Whole-body bioluminescent imaging of mice after intravenous injection with indicated plasmids (pNLuc, 5 μg/mice; pMEFI, 10 μg/mice) encapsulated within cationic lipids. (n =3) (E) Ex vivo organ bioluminescence from mice of (D) 72 hours after injection.

Effective systemic delivery of plasmid DNA presents a significant challenge^41^. We then evaluated MEFI’s efficacy in promoting plasmid DNA expression by means of in vivo and ex vivo luminescence. The indicated plasmids were encapsulated in a commercial cationic lipids. After 24 hours of intravenous injection, mice in the pNLuc alone group exhibited nearly undetectable expression of NLuc, whereas in the presence of MEFI, NLuc expression was ∼ 360-fold compared to that of the pNLuc alone (Figure 2D). A strong expression of NLuc persisted at least 72 hours facilitated by MEFI, and then the mice were sacrificed for organs ex vivo imaging. Weak NLuc expression was detected only in the lungs and spleens of mice treated with pNLuc alone, whereas the presence of MEFI resulted in substantially higher NLuc expression in most major organs compared to that of pNLuc alone: 501-fold higher in the lung, 147-fold higher in the liver, 85-fold higher in the spleen, 53-fold higher in the brain (Figure 2E), 27-fold higher in the kidneys, 50-fold higher in the heart, 35-fold higher in the pancreas, and 142-fold higher in the inguinal lymph nodes (Figure S2F). Importantly, the obvious expression of NLuc in the brain indicates that MEFI could transfer cargo mRNA across the blood-brain barrier (Figure 2E).

### MEFI enables robust and dureble expression of naked plasmids and interorgan cargo transfer *in vivo*

Intramuscular injection of naked plasmids is the one of most convenient methods of gene delivery, however, the effect is poor due to the low level of cargo protein expression^42,43^. Next, we attempt to evaluate whether MEFI can enhance cargo expression through intramuscular injection of naked plasmids. For the relative uniformity of the comparison, the same mouse was injected with pNLuc plus pGFP into the left caudal thigh muscle, while pNLuc plus pMEFI were injected into the right caudal thigh muscle. Indeed, we did not detect obvious NLuc expression from the left side, but a robust NLuc expression was observed on the right side up to two months post-injection. Moreover, a second administration still evoked significant and durable NLuc expression, suggesting that MEFI and its protein did not elicit an obvious adaptive immune response (Figure 3A-B). We then examined the ability of MEFI to deliver therapeutic naked pDNA cargo via intramuscular injection. 28 days post-administration, we detected therapeutic levels of human erythropoietin (hEPO) in the serum of mice injected with phEPO/pMEFI, whereas no detectable hEPO was present in the absence of MEFI (Figure 3C).

**Figure 3.**
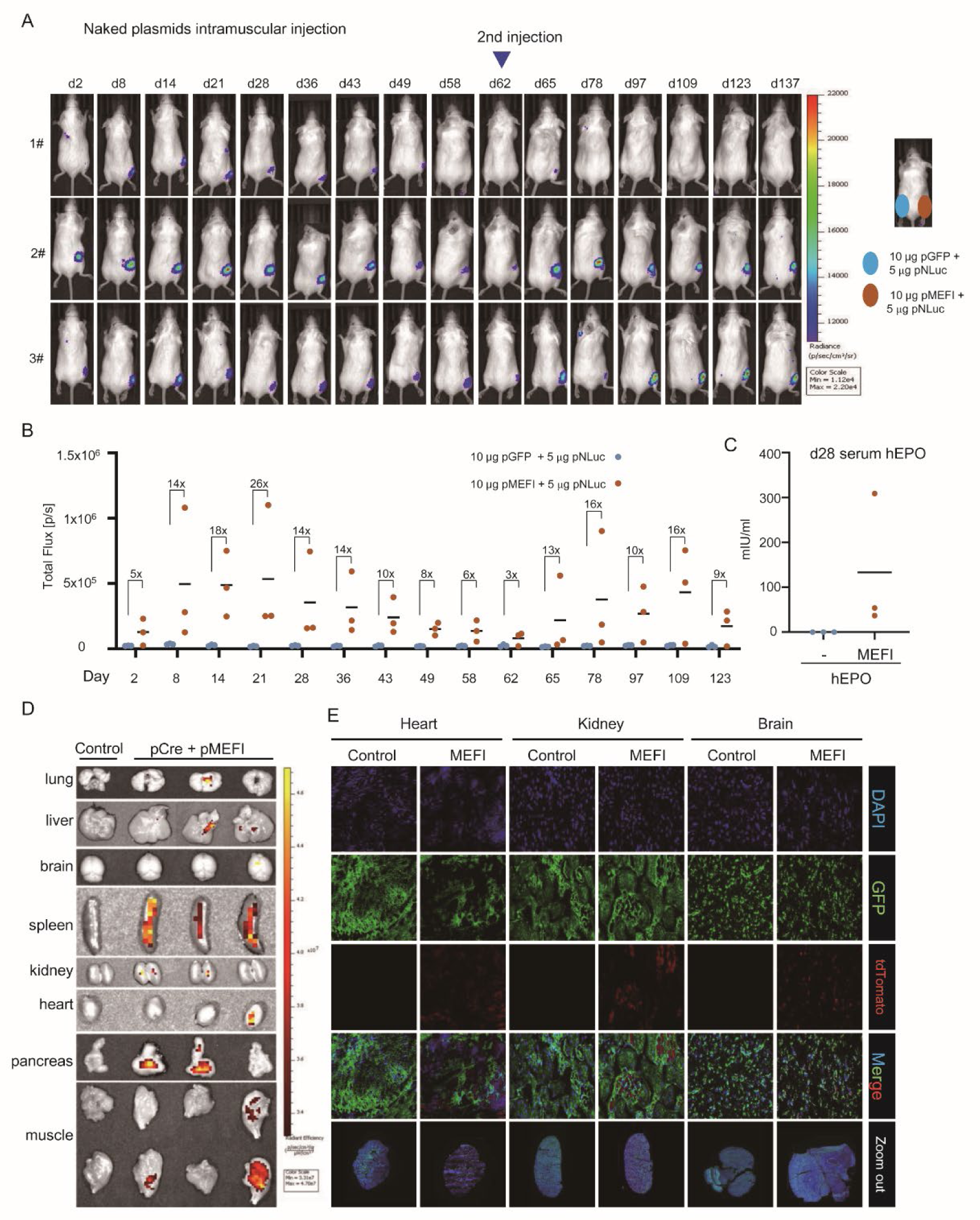
MEFI enables robust and dureble expression of naked plasmids and interorgan cargo transfer in vivo. (A) Whole-body bioluminescent imaging of mice after intramuscular injection with indicated naked plasmids. The same mouse was injected with pNLuc plus pGFP into the left caudal thigh muscle, while pNLuc plus pMEFI were injected into the right caudal thigh muscle. (n =3) (B) Quantification of bioluminescent signal from (A). (C) Serum hEPO concentrations in mice which were intramuscular injected with indicated naked plasmids 28 days ago, n = 3 biologically independent mice per group. (D) Ex vivo organ tdTomato fluorescence from mice that were intramuscularly injected with the indicated naked plasmids 7 days prior (n=3). (E) Confocal imaging of tissue sections of mice from (D).

To further confirm whether MEFI can mediate cargo mRNA transfer *in vivo*, we utilized genetically engineered ZsGreen/tdTomato conversion reporter mice (CAG-LoxP-ZsGreen-Stop-LoxP-tdTomato) containing a LoxP-flanked ZsGreen-STOP cassette that initially expresses ZsGreen but prevents the expression of the tdTomato protein. Once the ZsGreen-STOP cassette is deleted by Cre, ZsGreen expression is turned off, and tdTomato fluorescence is turned on, allowing for the detection of Cre-mediated gene-edited cells. Seven days following the intramuscular injection of the MS2-tagged Cre plasmid and pMEFI, tdTomato fluorescence was readily detected in some organs through ex vivo imaging (Figure 3D). Additionally, tdTomato-positive cells were easily observed using confocal imaging of tissue sections (Figures 3E, S3B). Specifically, tdTomato-positive cells were emerging in the brain, demonstrating that MEFI facilitates Cre mRNA crossing the blood-brain barrier (Figure 3E). These findings suggest that MEFI can facilitate interorgan cargo mRNA transfer in mice.

### Cell type-specific cytotoxicity conducted by scFv-MEFI

Next, we aim to modify MEFI for cell-specific delivery. Reports indicate that by fusing a single-chain variable fragment (scFv) between the secretion signal and transmembrane domain of VSVg, its antigen targeting tropism can be achieved^44–46^. Inspired by this, we constructed MEFI variants targeting CD19 and tyrosinase-related protein 1 (TYRP1, a melanoma surface antigen) by replacing the extracellular domain of VSVg with the corresponding anti-CD19 or anti-TYRP1 scFv, respectively (Figure 4A). Then, we used NLuc as cargo to verify the specific targeting of scFv-MEFI. The chemiluminescence assay revealed that in the presence of MEFI or anti-CD19 scFv-MEFI, the expression of NLuc in Naml6 cells (CD19^+^), were almost equal. However, the anti-CD19 scFv-MEFI was inefficient in transferring NLuc mRNA to K562 cells (CD19^-^) compared to native MEFI (Figure 4B). A similar result was obtained in the items of anti-TYRP1 MEFI in TYRP1^+^ B16 and TYRP1^-^ GL261 cells (Figure 4C)

**Figure 4.**
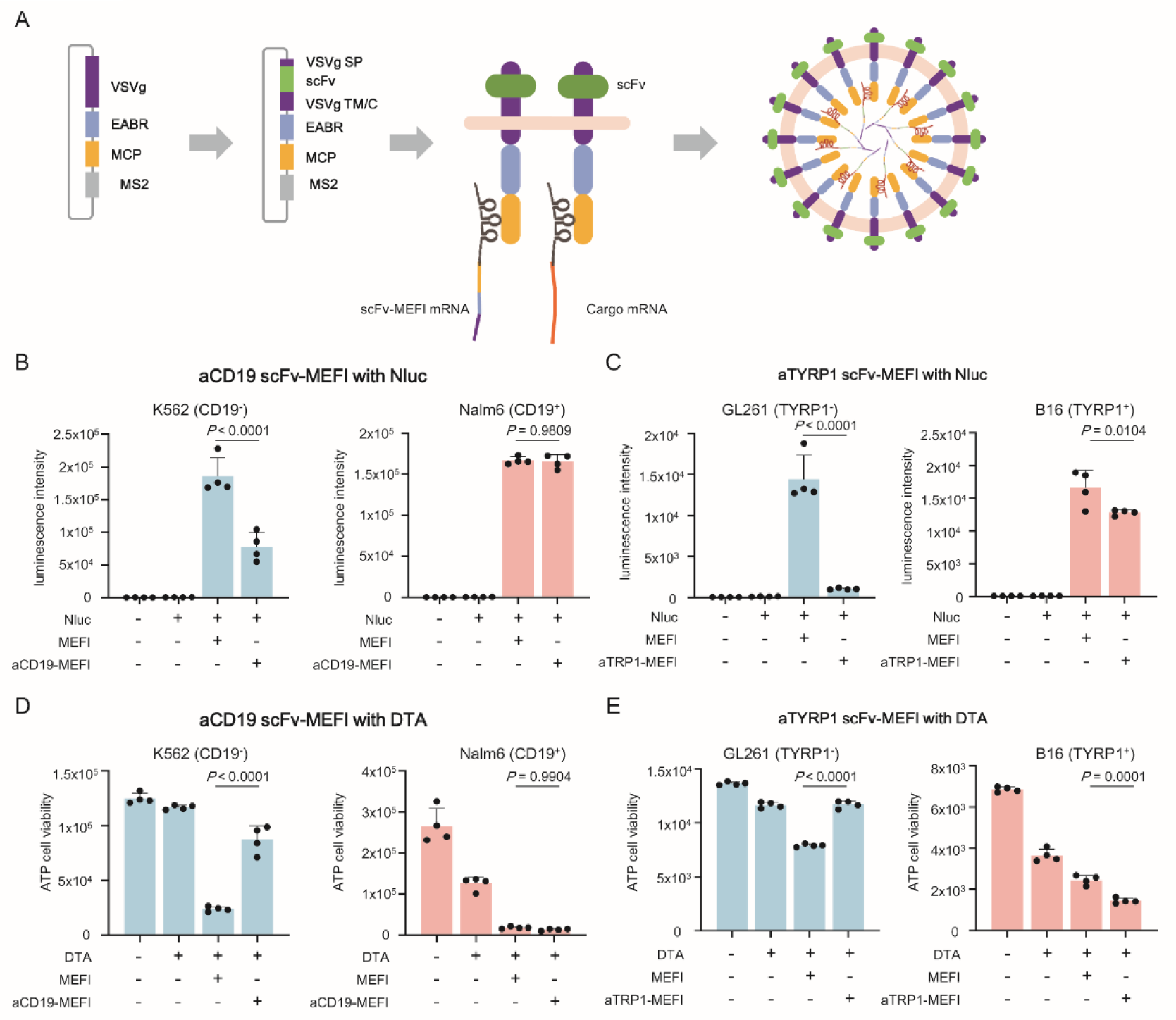
Cell type-specific cytotoxicity conducted by scFv-MEFI. (A) The schematic diagram shows scFv-MEFI facilitates the assembly of VLP, which packages both cargo and itself mRNA. (B-C) The indicated plasmids were transfected into 293T cell. After 36 hours, the supernatants were filtered and added to indicated cells. 24 hours later, the NLuc expression was measured using chemiluminescence. (D-E) The indicated plasmids were transfected into 293T cell. After 36 hours, the supernatants were filtered and added to indicated cells. 24 hours later, the cell viablity was measured by ATP assay.

Diphtheria toxin A chain (DTA), a widely used inducer of cell death^47^, was selected as the cargo to test the cell type specific killing of scFv-MEFI. Twenty-four hours after adding the indicated transiently co-transfected supernatant, the ATP assay demonstrated the cell viability of Nalm6 cells were both decreased in the present of MEFI and anti-CD19 scFv-MEFI. However, compared with that of anti-CD19 MEFI, only the MEFI DTA-treated K562 cells were obviously killed (Figure 4D). A similar result was obtained in the items of anti-TYRP1 MEFI with TYRP1^+^ B16 and TYRP1^-^ GL261 cells, confirming its cell-type cytotoxicity (Figure 4E).

Compared to traditional small-molecule proteolysis targeting chimeras (PROTACs), biological PROTACs expand the range of targeted proteins with a more flexible E3 ligase selection^48–50^. However, the challenges associated with specific intracellular delivery limit their application. We thus utilized a bioPROTAC, targeting the oncology-related protein Proliferating Cell Nuclear Antigen (PCNA), as a cargo to assess the specific delivery of scFv-MEFI. The ATP assay revealed a more significant reduction in the viability of Nalm6 cells treated with the supernatant from anti-CD19-MEFI/aPCNA-PROTAC compared to those treated with the supernatant from MEFI/aPCNA-PROTAC. However, the cell viability of K562 cells was more decreased in the presence of MEFI-PROTAC than with anti-CD19 scFv-MEFI-PROTAC (Figure S4A). A similar result was obtained for the anti-TYRP1 MEFI with both TYRP1^+^ B16 and TYRP1^-^ GL261 cells (Figure S4B). These results demonstrate that MEFI can be modified for cell-specific targeting to deliver diverse therapeutic cargoes.

## Discussion

The development of the mRNA Exporting and Ferrying Implement (MEFI) represents a paradigm shift in mRNA therapeutics, as it enables iterative, cell-to-cell transfer of cargo mRNA. Unlike conventional mRNA delivery systems where mRNA functions transiently within a single cell, MEFI harnesses a self-replicating VLP system to amplify cargo expression across cellular barriers, significantly enhancing therapeutic potency and durability.

MEFI’s core innovation lies in its ability to package both cargo mRNA and its own mRNA into fusogenic VLPs, enabling continuous propagation from primary to secondary target cells. This "multiple-use" effect addresses a fundamental limitation of short-lived expression of current mRNA therapies by extending cargo availability across cell populations. MEFI enhances low-efficiency delivery methods (e.g., naked plasmid intramuscular injection) by orders of magnitude, potentially revolutionizing DNA vaccine strategies. The sustained protein production by naked plasmid intramuscular injection for two months underscore its potential for long-term treatments, such as protein replacement therapies. The ability to cross the blood-brain barrier further expands applications to neurological disorders. By replacing VSVg’s extracellular domain with specific scFvs (anti-CD19/anti-TYRP1), MEFI achieves cell-type-specific cargo delivery. This modular targeting system enabled selective cytotoxicity in antigen-positive cells using lethal cargoes (DTA) or bioPROTACs, while sparing off-target cells. Such precision could mitigate systemic toxicity in oncology and expand the scope of biologics delivery, including previously undruggable intracellular targets.

Unlike conventional eVLPs requiring multiple viral components, MEFI’s single VEM fusion protein simplifies production and reduces immunogenicity. Its capsid-independent assembly bypasses packaging constraints of lentiviral systems. While delivered mRNA to primary cells, MEFI enables secondary spread, amplifying effects in poorly accessible tissues (e.g., brain, solid tumors). Although adaptive immunity against MEFI was minimal, long-term studies must assess anti-fusogen responses. Engineering hypoimmunogenic variants (e.g., humanized fusogens) could mitigate risks. Attenuation over transfer rounds may reflect mRNA degradation or translational silencing. Incorporating RNA-stabilizing elements (e.g., UTR optimization, nucleotide modifications or circular mRNA) could enhance persistence. While scFv-MEFI showed promising specificity, in vivo validation in complex microenvironments is needed. Alternative targeting moieties (e.g., de novo designed binders^52^) may improve affinity. Expanding beyond MS2-MCP systems (e.g., using L7Ae-kink-turn^53^) could enable alternative applications.

MEFI establishes a modular, extensible platform that transforms mRNA from a transient payload into a self-propagating therapeutic agent. Its capacity for iterative delivery, systemic amplification, and cell-specific targeting positions it as a platform technology for next-generation biologics. Future work will focus on optimizing safety profiles, extending cargo diversity (e.g., base editors, transcription factors, large transgenes), and advancing toward clinical translation in oncology, genetic disorders, and regenerative medicine in animal models, organoids and patients.

## RESOURCE AVAILABILITY

### Lead contact

Further information and requests for resources and reagents should be directed to and will be fulfilled by the lead contact, Xiushan Yin

## DATA AVAILABILITY STATEMENT

The data that support the findings of this study are available from the corresponding author upon reasonable request.

## ACKNOWLEDGMENTS

This work was supported by Roc Rock Biotechnology (Suzhou) Co., Ltd. The research project was funded partially by the the National Natural Science Foundation of China (32371490) and Beijing Municipal Natural Science Foundation (5252022) to L.Z. Additional support was provided by the 2023 National Center for Biotechnology Innovation in Cellular Therapy Technology Research Project (2023XB0010) to X.Y.

## AUTHOR CONTRIBUTIONS

L.Z. and X.S.Y. conceptualized this work. L.Z., X.S.Y., Y.G., H.L., Y.X., Y.L., Y.F.L., S.L., X.J.S., X.B.F., J.H.L., Q.Y.Z., and X.S.Y. developed the methodology. Y.G., L.Z., X.S.Y., H.L., Y.X., Y.L., Y.F.L., S.L., X.J.S., X.B.F., J.H.L. and Q.Y.Z. carried out the investigation. L.Z., Y.G., H.L., Y.X., Y.L., Y.F.L., S.L., X.J.S., X.B.F. and J.H.L. created visual representations. L.Z., Y.G. and X.S.Y. wrote the initial draft and L.Z., Y.G., X.S.Y., H.L., Y.X. and X.B.F. reviewed and edited the manuscript. L.Z. and X.S.Y. were responsible for funding acquisition and project administration.

## KEY RESOURCES TABLE

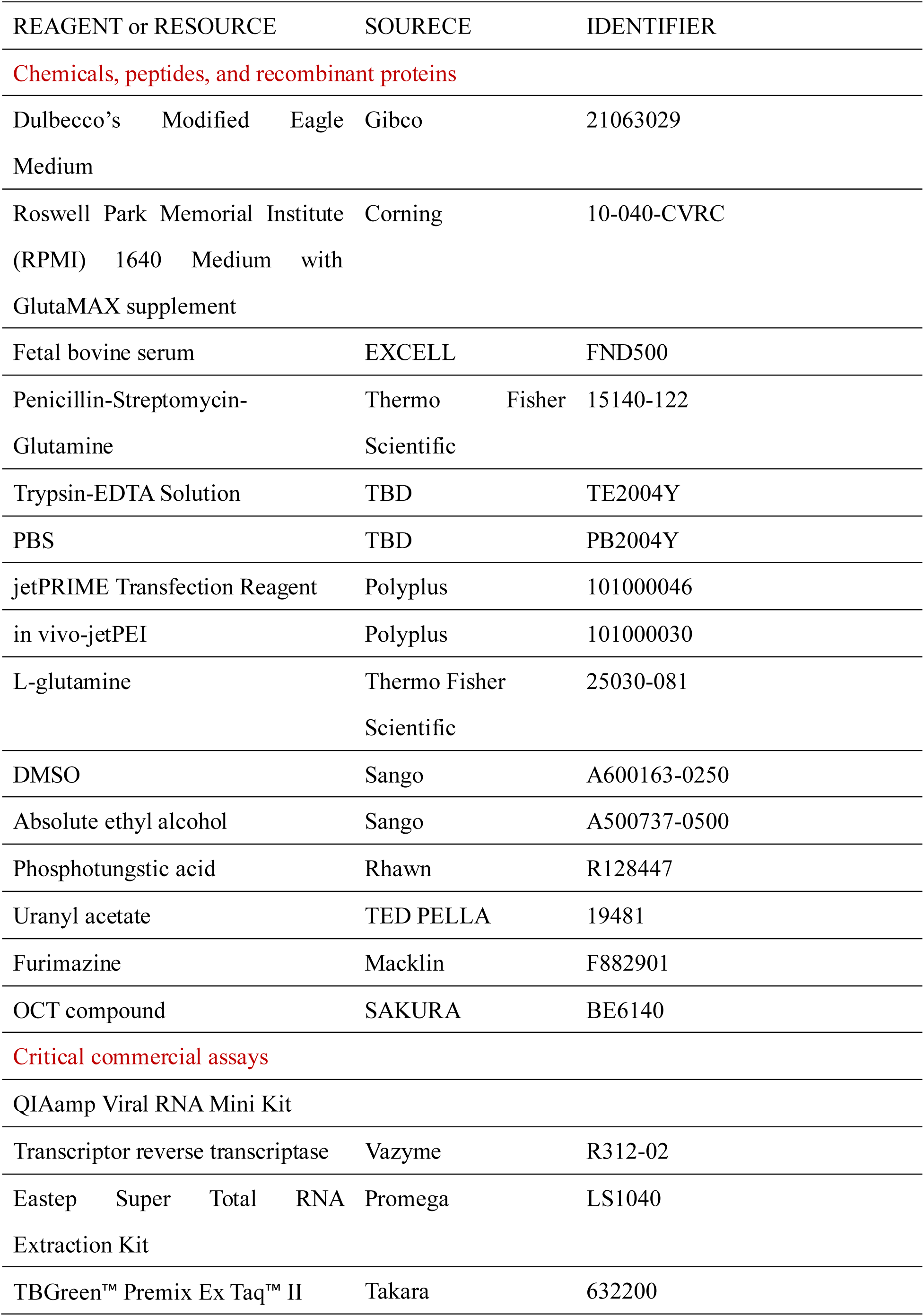
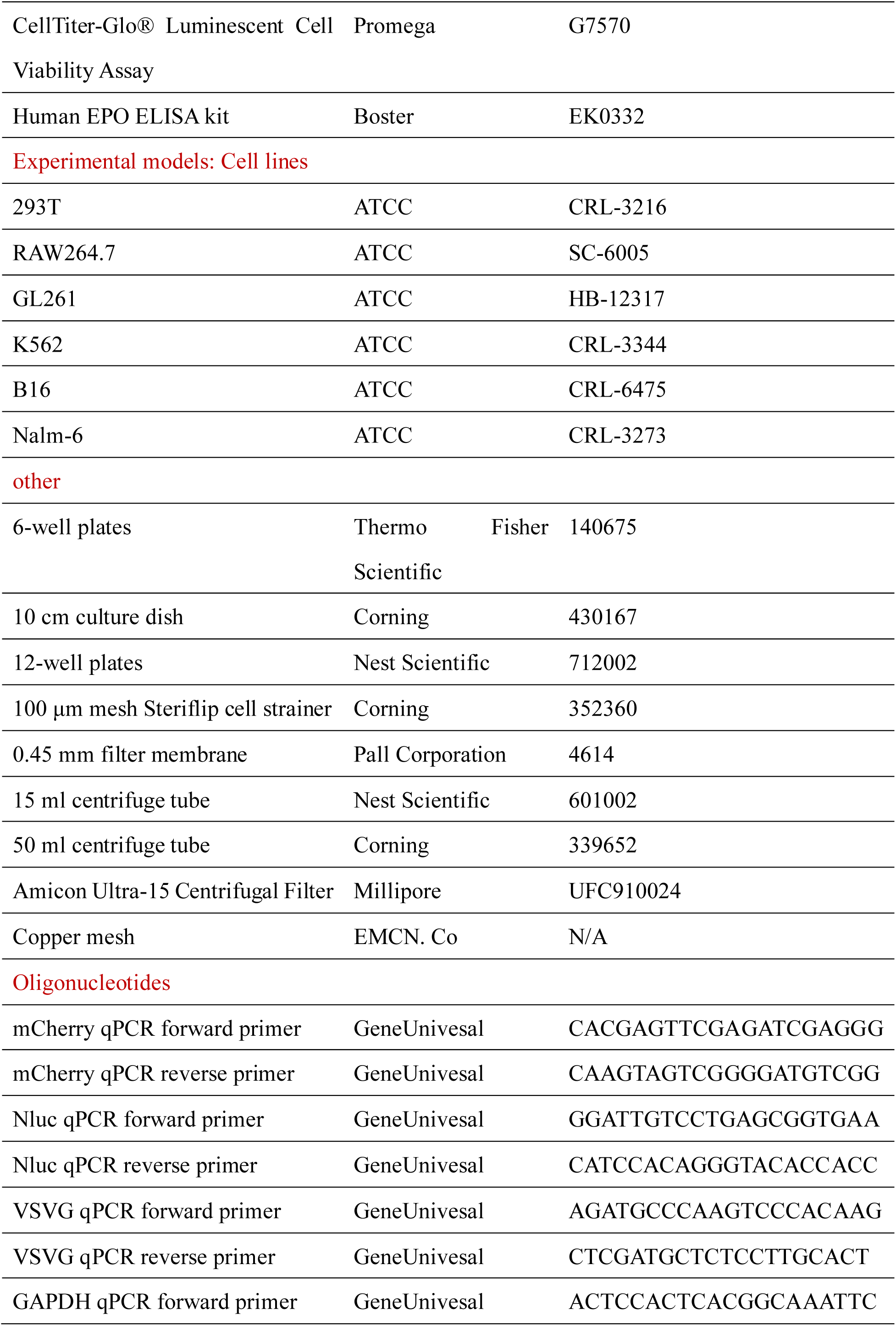
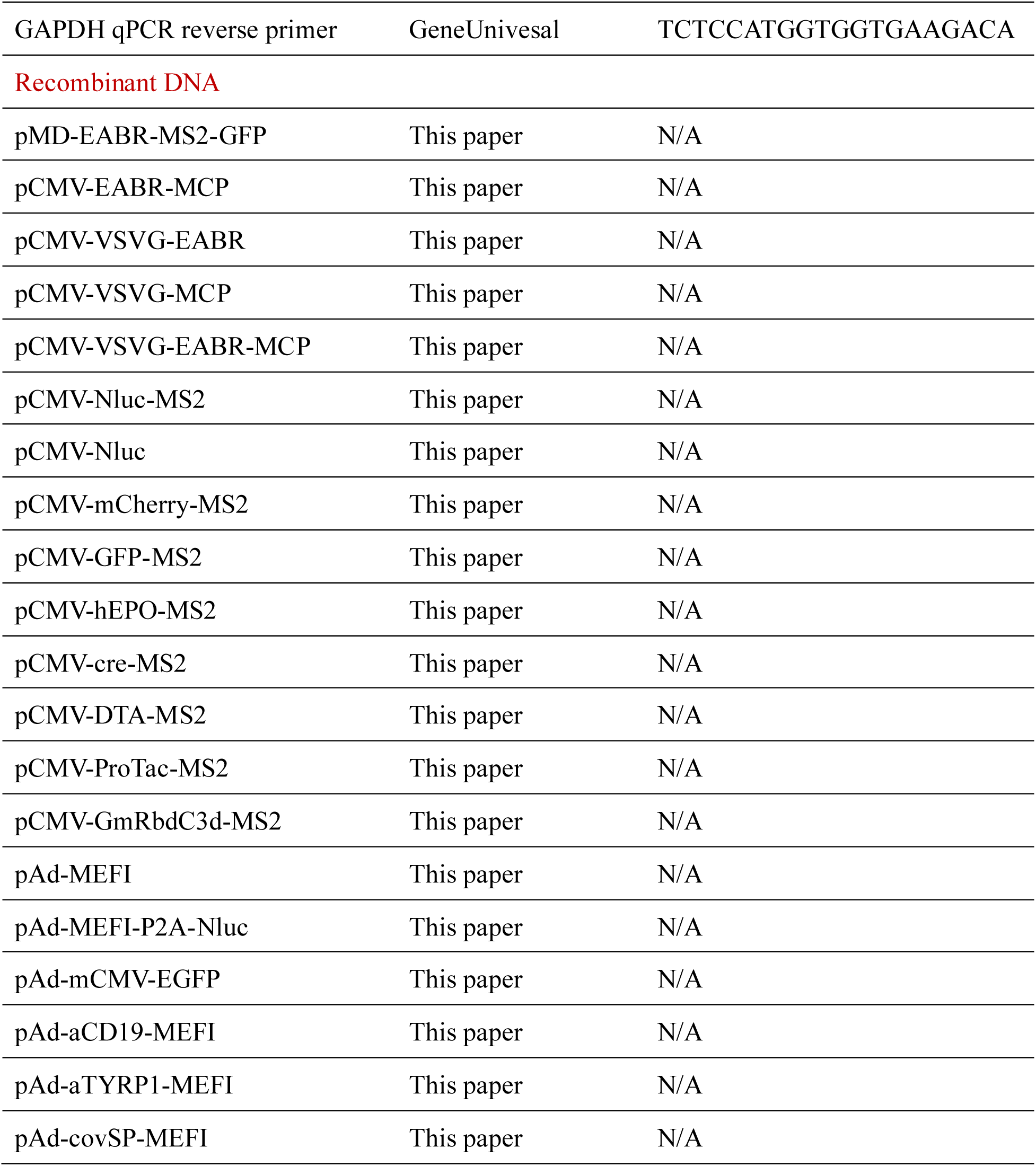

## EXPERIMENTAL MODEL AND STUDY PARTICIPANT DETAILS

### Cell lines

HEK293T and GL261 cells were cultured in Dulbecco’s modified Eagle’s medium (DMEM, Gibco) supplemented with 10% heat-inactivated fetal bovine serum (FBS, EXCELL) and 1% penicillin-streptomycin (Gibco) at 37℃ and 5% CO_2_. RAW264.7, K562, Nalm-6, and B16 cells are cultured in Roswell Park Memorial Institute (RPMI) 1640 Medium supplemented with 10% FBS and 1% penicillin-streptomycin.

### Plasmid construction

Constructs used in this study are listed in the key resources table. Some constructs were generated by standard cloning procedures, in which inserts and linearized backbones were generated by polymerase chain reaction (PCR) or restriction digest. The remaining constructs were designed by the authors and synthesized by GeneUnivesal.

### Mouse studies

Animals were group-housed in IVCs under SPF conditions, with constant temperature and humidity with lighting on a fixed light/dark cycle (12-hours/12-hours). 5 – 7 weeks old homozygous mouse littermates were randomly assigned to experimental groups. No mice were involved in previous procedures.

In vivo studies were done using adult, 8–10-week-old, male BALB/c (Charles River Laboratories, Beijing, China) (Figure 2D). Intravenous injection occurred via the lateral tail vein with 100 μL of the test agent. Furimazine solution was injected intraperitoneally at the specified time point into both acquisition and analysis of optical images.

In vivo studies were done using adult, 8–10-week-old, female BALB/c (Charles River Laboratories, BeiJing, China) (Figure 3A). Intramuscular injection occurred in the lateral muscle group of the thigh with 80 μL of the test agent.

In vivo studies were done using adult, 8–12-week-old, male B6-G/R (RocRock Laboratories, Shenyang, China) (Figure 3D). Intramuscular injection occurred in the lateral muscle group of the thigh with 80 μL of the test agent.

## METHODS DETAILS

### VLP production

VLPs in our paper were produced by transient transfection of HEK293T cells. HEK293T cells were seeded in 6-well plates at a density of 5 x 10^5^ cells per well. After 24h, cells were transfected using the jetPRIME transfection reagent according to the manufacturer’s protocols. The reagent volume to DNA mass ratio(V/m) was 1:2 (fill-up with JetOPTIMUS Buffer, 10 min incubation at room temperature for complex formation). The detailed plasmid stoichiometries of the different systems are shown in this paper. The media was changed after 6 hours, and VLP supernatant was harvested at 36 hr and centrifuged for 5 min at 500 g to remove cell debris. The clarified VLP-containing supernatant was filtered through a 0.45-mm filter. For VLPs that were used in cell culture, unless otherwise stated, the filtered supernatant was concentrated 100-fold using Amicon Ultra-15 Centrifugal Filter. Ultracentrifugation was performed at 135,000 g for 2 h (4℃) using an SW 40Ti rotor in an Optima XPN-80 Ultracentrifuge (Beckman Coulter). Following ultracentrifugation, VLP pellets were resuspended in cold PBS. Samples were aliquoted and stored at -20℃.

For making specific receptor targeting and integrating VLPs, VSV-G mutant was used instead.

### Lentiviral infection and viral incubation assay

To evaluate VLPs, 20 μL concentrated EABR-VLP was added into RAW264.7 cells in 12 well plate. 2 days later, GFP signal was measured by flow cytometry. To define the roles of VSVg, EABR, and MCP in mRNA packaging and delivery, 40 μL concentrated VLPs were added into RAW264.7 cells in 6 well plate. Upon 2 days incubation, cells were subjected to flow cytometry to quantify mCherry+ cells.

### Negative-stain transmission electron microscopy

VLPs were centrifuged for 5 min at 500 g to remove debris and then the clarified supernatant were ultracentrifuged for 2 hrs at 135,000 g. A drop of 10 μL of sample was applied to a carbon coated grid that had been glow discharged for 1 min in air, and the grids were immediately negatively stained using 2% phosphotungstic acid for 60 s. Grids were examined in a H-7800 operated at 80-120 kV. Images were taken using HITACHI Transmission Electron Microscope.

### Bioluminescence quantification

NLuc signal was read on-plate 48 hours post-transfection. Measurements were taken on a BioTek synergy H1 plate reads with 1 s acquisition time right after the addition of Furimazine (Macklin) for NLuc.

### Flow cytometry

48 h post-incubation by VLPs, cells were gently detached in a suitable volume of accutase, pelleted (500 g, 5 min), and resuspended in PBS. Analysis was performed on the BD FACS machine. Data were analyzed with FlowJo. The main population of the cells was gated according to their FSC-A and SSC-A. About the single cells were gated using FSC-A and FSC-H.

### Reverse transcription and quantitative PCR

We used an RT-qPCR assay to measure the abundance of specific RNA molecules in VLP or cellular RNA. The VLP RNA was extracted from 10 mL of supernatant using the Viral RNA Mini kit according to manufacturer’s instructions. The cellular RNA was extracted from cells using the Eastep Super Total RNA Extraction Kit according to manufacturer’s instructions. RNA was then reverse transcribed using Transcriptor reverse transcriptase mix at 42 ℃ for 15 mins, and 95 ℃ for 3 mins. Quantitative polymerase chain reaction (qPCR) was performed on the CFX96 Touch system (Biorad) using SYBR Green Supermix with reverse transcription product as input, final concentration of 400 nM per primer, and thermal cycling profile consisting of 95 ℃ for 30 seconds, followed by 40 cycles of 95 ℃ for 10 seconds and 60 ℃ for 30 seconds. Primer sequences are listed in key resources table. Negative controls consisting of supernatant from HEK293T cells subjected to mock transfections without DNA.

### ELISAs

Serum EPO levels were quantified using the BOSTER Human EPO Assay kit following the manufacturer’s instructions. First of all, plates are coated with biotinylated capture antibody prior to sample and standard administration. Samples and standards are incubated on plate for 90 mins at 37 ℃, following which detection antibody is added for 1 hour. Avidin-peroxidase complex was added to the plate for 30 mins incubation, and then it was analyzed with the BioTek synergy H1.

### Cell viability assays

Cell viability was quantified using a Promega CellTiter-Glo luminescent cell viability kit. 5x10^4^ cells (forB16, Nalm-6, GL261 and K562) were seeded in 100 μL of media per well. The cells were allowed to adhere for 24 h before treatment with VLPs. After 48 h of VLP incubation, 100 μL of CellTiter-Glo reagent was added to each well in the dark. Cells were incubated for 10 min at room temperature and the 100 μL of solution was transferred into black 96-well flat bottom plates, and the luminescence was measured on a BioTek synergy H1 plate reads with 1 s acquisition time.

### Histofluorescence staining

Tissues were fixed in 4% paraformaldehyde (PFA) for 24 h, dehydrated in 30% sucrose solution. Frozen sections were preserved in OCT compound, frozen, sectioned at 3 µm, and stored at -80 °C. Prior to staining, all samples were rinsed with ddH2O. The sections were then incubated 5 minutes at room temperature with 5 μg/mL DAPI away from light, then washed with ddH2O. Images were captured using a confocal microscopy system (LSM 700, Zeiss, Jena, Germany). Image analysis was performed in ImageJ.

