## Supplemental Figure for "Iterative delivery of mRNA by MEFI"

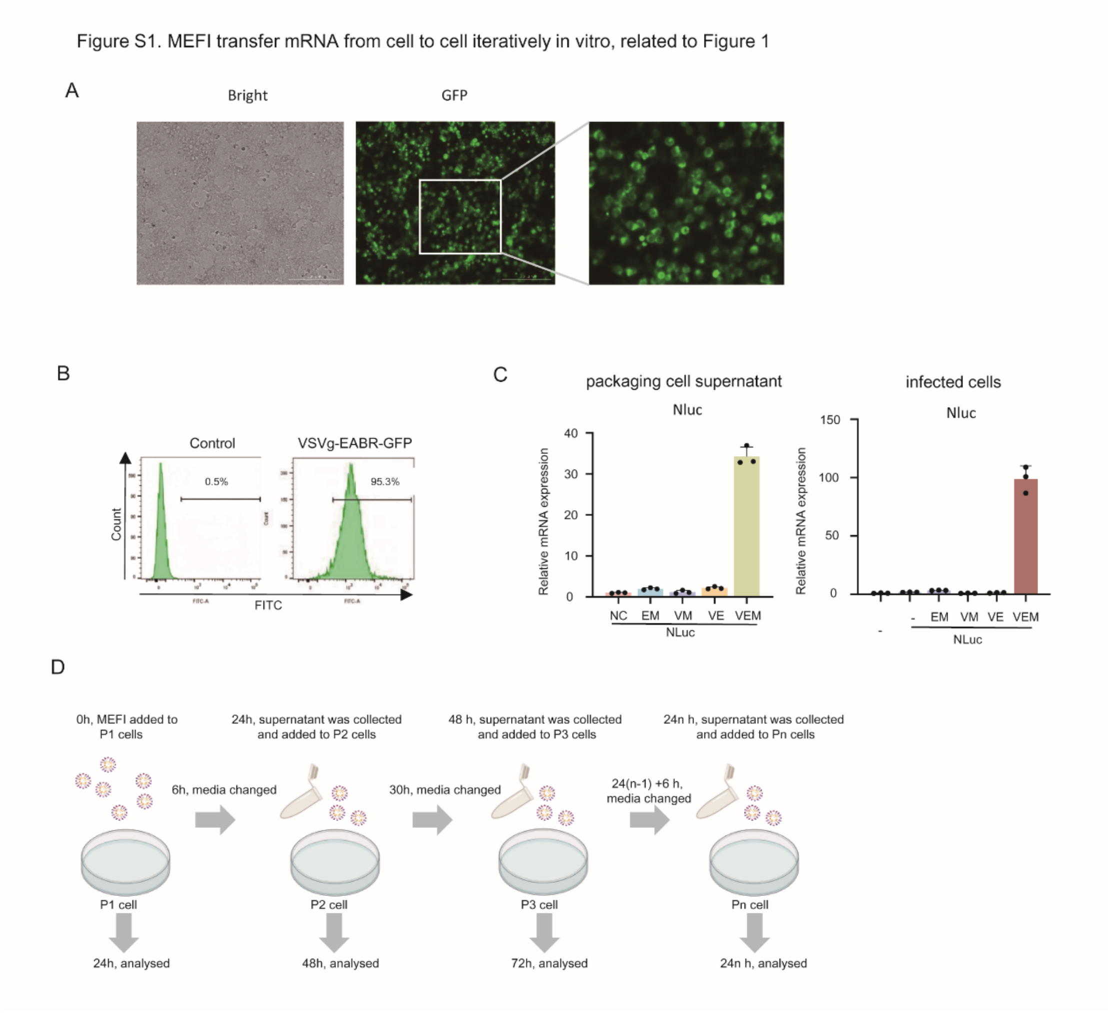


**Figure S1. MEFI encodes fusion protein packages mRNA into self-propagating VLPs, enabling continuous cell-to-cell mRNA transfer, related to Figure 1**

(A) 293T cells were tranfected with VSVg-EABR-GFP plasmid for 24 hours before fluorescence images were taken.

(B) Flow cytometry conducted on cells from Fig1 C

(C) The relative quantity of mCherry-MS2 mRNA in RAW264.7 cells infected with the supernatants indicated for 24 hours by qPCR assay. Each dot represents one technical replicate.

(D) The flow diagram of iterative infection related to Figure 1M

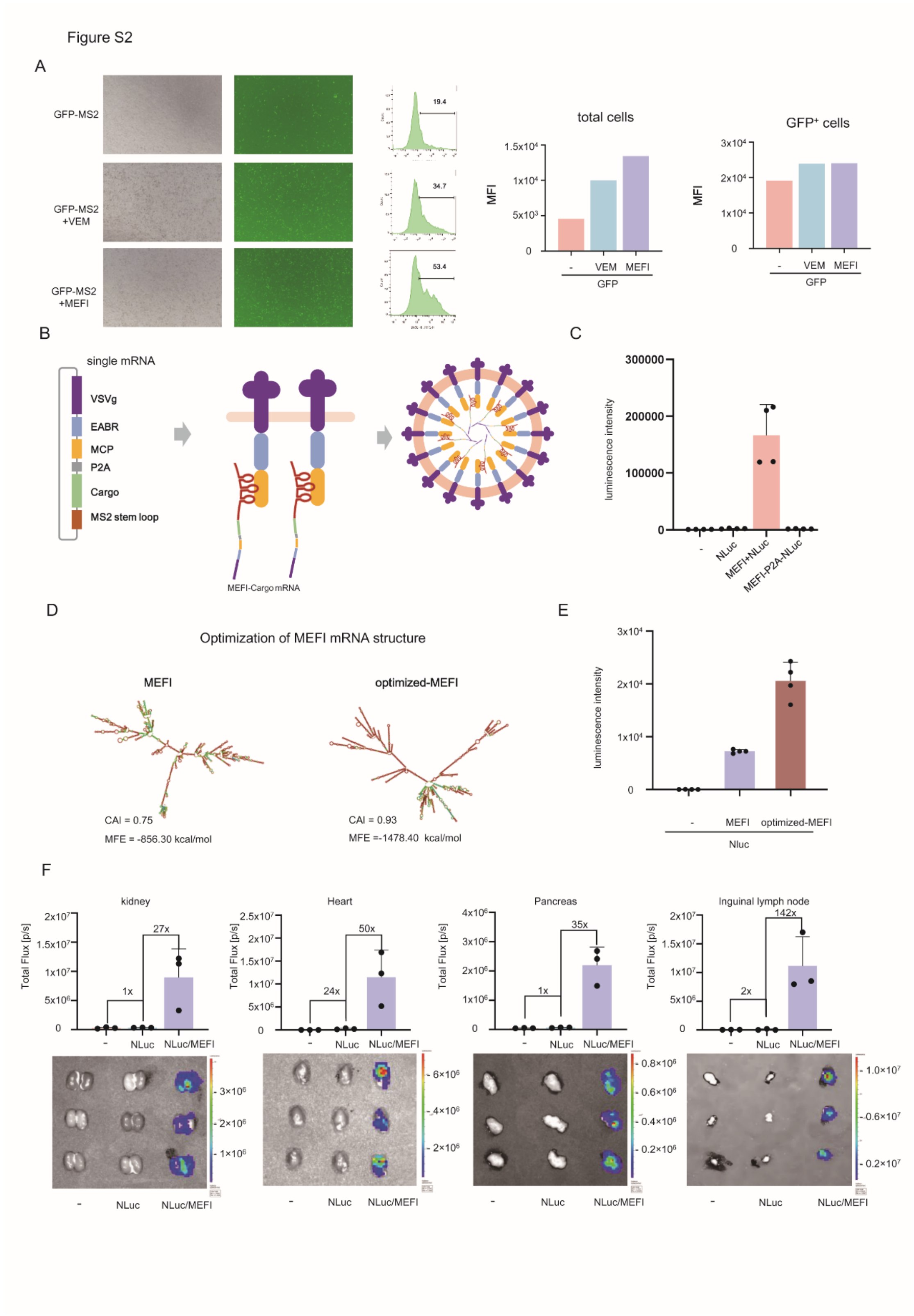


**Figure S2. The optimized MEFI system enhances cargo expression in vitro and in vivo, related to Figure 2**

(A) 293T cells were tranfected with indicated plasmids for 36 hours. Then, fluorescence images were taken and flow cytometry was carried out for GFP.

(B) The schematic diagram shows single Cargo-MEFI mRNA system.

(C) The indicated plasmids were transfected into 293T cells. After 36 hours, the supernatants were filtered and added to RAW264.7 cells. 24 hours later, the NLuc expression was measured using chemiluminescence.

(D) The secondary structures of MEFI (left) and optimally MEFI (right) mRNAs.

(E) The indicated plasmids were transfected into RAW264.7 cell. After 36 hours, the NLuc expression was measured using chemiluminescence.

(F) Ex vivo organ bioluminescence from mice of (Figure 2D) 72 hours after injection.

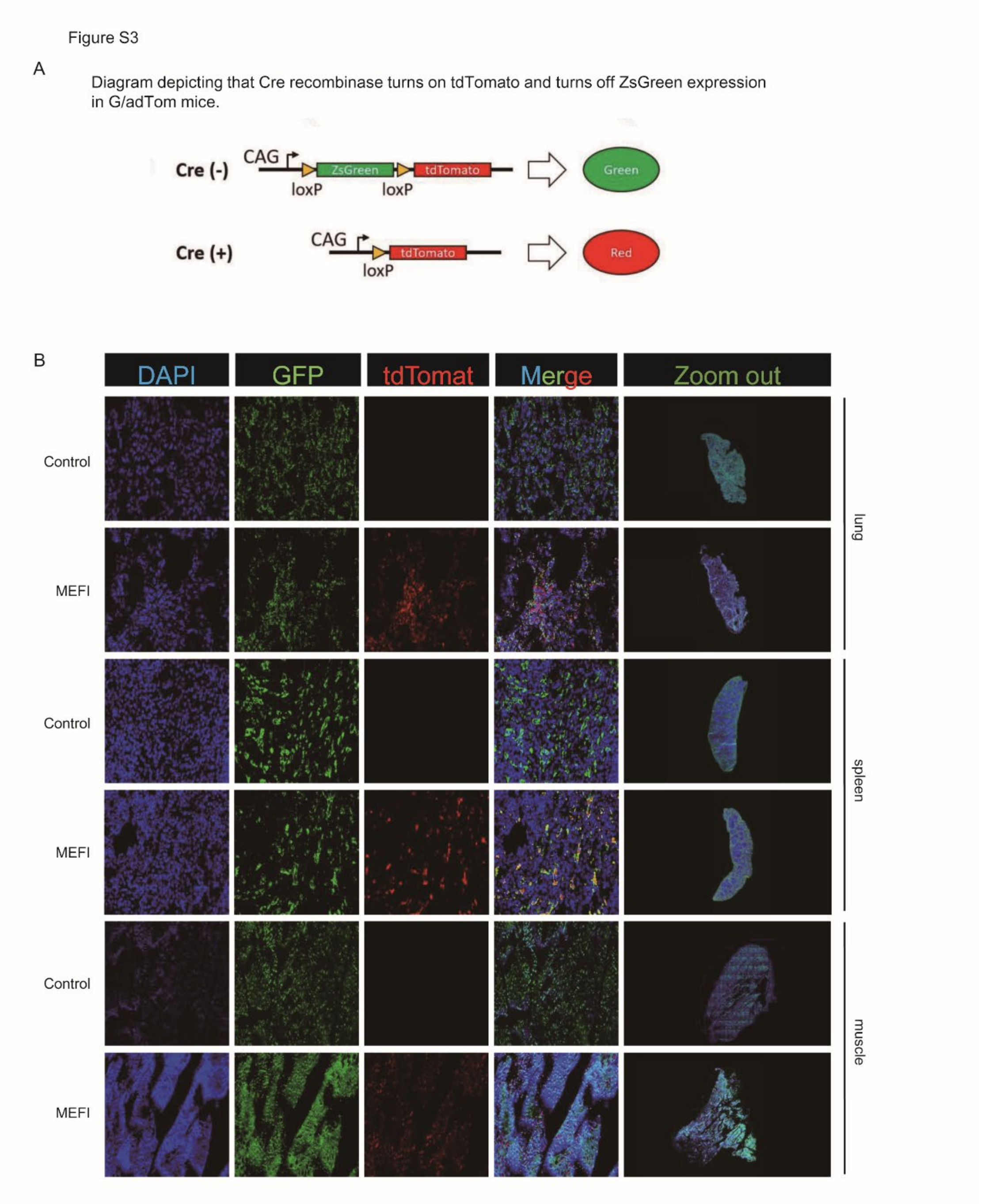


**Figure S3. MEFI enables robust and dureble expression of naked plasmids and interorgan cargo transfer in vivo, related to Figure 3.**

(A) Scheme illustrating that Cre recombinase turns on tdTomato and turns off ZsGreen expression in G/adTom mice.

(B) Confocal imaging of tissue sections of mice from (Figure 3D).

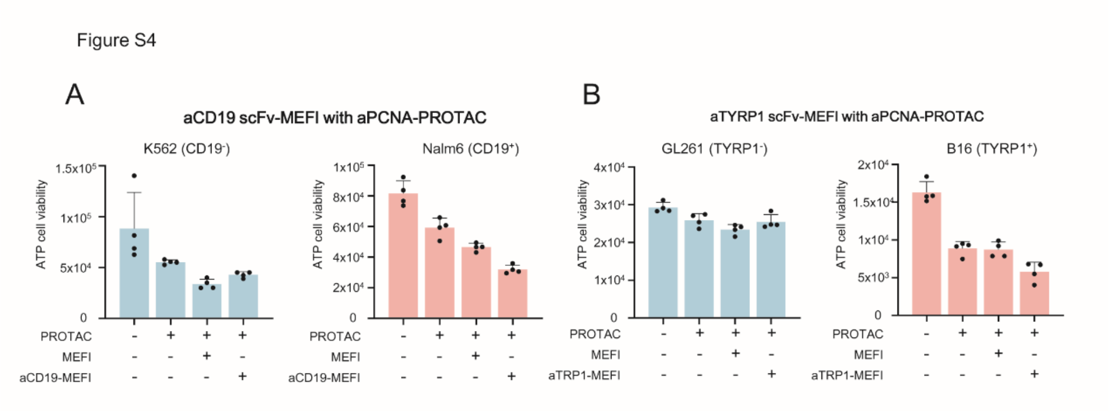


**Figure S4. Cell type-specific cytotoxicity conducted by scFv-MEFI**

(A) The indicated plasmids were transfected into 293T cells. After 36 hours, the supernatants were filtered and added to indicated cells. 24 hours later, the cell viablity was measured by ATP assay.
